# Short Tandem Repeats of Human Genome Are Intrinsically Unstable in Cultured Cells *in vivo*

**DOI:** 10.1101/2023.02.12.528158

**Authors:** Yuzhe Liu, Jinhuan Li, Qiang Wu

**Affiliations:** Center for Comparative Biomedicine, Ministry of Education Key Laboratory of Systems Biomedicine, State Key Laboratory of Oncogenes and Related Genes, Institute of Systems Biomedicine, Shanghai Jiao Tong University, Shanghai 200240, China; WLA Laboratory, Shanghai 201203, China

## Abstract

Short tandem repeats (STRs) are a class of abundant structural or functional elements in the human genome and exhibit a polymorphic nature of repeat length and genetic variation within human populations. Interestingly, STR expansions underlie about 60 neurological disorders. However, “stutter” artifacts or noises render it difficult to investigate the pathogenesis of STR expansions. Here, we systematically investigated STR instability in cultured human cells using GC-rich CAG and AT-rich ATTCT tandem repeats as examples. We found that triplicate bidirectional Sanger sequencing with PCR amplification under proper conditions can reliably assess STR length. In addition, we found that next-generation sequencing with paired-end reads bidirectionally covering STR regions can accurately and reliably assay STR length. Finally, we found that STRs are intrinsically unstable in cultured human cell populations and during single-cell cloning. Our data suggest a general method for accurately and reliably assessing STR length and have important implications in investigating pathogenesis of STR expansion diseases.

## Introduction

In eukaryotes, repetitive DNA sequences are abundant and widely distributed throughout their genomes (Treangen and Salzberg, 2012; Balzano et al., 2021). In particular, approximately 3% of the human genome comprises short tandem repeats (STRs, also known as microsatellites) usually with small units [typically 2-12 base pairs (bp)] (Gymrek, 2017; Malik et al., 2021). STRs have intrinsic instabilities of the nucleotide repeat tracts and display a high degree of length polymorphisms and variations within human populations (Balzano et al., 2021). Interestingly, expansions of a subset of these nucleotide repeat tracts underlie about 60 neurological disorders, including amyotrophic lateral sclerosis (ALS), fragile X syndrome (FXS), Huntington’s disease (HD), myotonic dystrophy (DM), spinocerebellar ataxias 10 and 12 (SCA10 and SCA12) (Holmes et al., 1999; Matsuura et al., 2000; La Spada and Taylor, 2010; Gall-Duncan et al., 2022). Specifically, most of these disorders result from the expansions of (CAG/CTG)_n_, (CGG)_n_, (GAA)_n_, (CCTG)_n_, (ATTCT)_n_, and even the dodecanucleotide repeats of (C_4_GC_4_GCG)_n_ (Mirkin, 2007; Gall-Duncan et al., 2022). According to the GC content, these STRs can be divided into GC-rich STRs and AT-rich STRs (Schroder et al., 2022). Intriguingly, nearly all neurological disease-associated STRs are located at the boundaries of topologically associating domains (TADs) (Sun et al., 2018), suggesting their central roles in 3D genome architecture. Finally, STR instability is a general feature of many types of cancers and accurate STR genotyping is central for forensic science (Page and Graham, 2008).

A CAG/CTG repeat expansion from 7-45 copies to 55-78 copies in the 5’UTR region of the *PPP2R2B* (Protein Phosphatase 2 Regulatory Subunit Bβ) gene was found to induce autosomal dominant SCA12 with action tremor, degeneration of the cerebellum, and spinocerebellar ataxia (Holmes et al., 1999; Gatchel and Zoghbi, 2005; Dong et al., 2015). In addition, a CAG repeat expansion encoding a polyglutamine (polyQ) stretch of the androgen receptor underlies spinal and bulbar muscular atrophy (SBMA) (La spada et al., 1991). Finally, expansion of pentanucleotide (ATTCT) repeats in the intron 9 of the *ATXN10* (Ataxin 10) gene from ∼10-22 copies to ∼850-4500 copies leads to a genetic disease associated with spinocerebellar ataxia and epilepsy, known as SCA10 (Matsuura et al., 2000; Guo and Lam, 2020).

One difficulty in the investigation of pathogenesis of STR expansion diseases is the establishment of accurate methods for detecting STR length or exact numbers of tandem repeat units. This is not trivial because of the so-called “stutter” artifacts in sequencing traces and “shadow” bands in agarose gels (Fig. S1A). Stutter artifacts often exhibit one or several repeat units shorter or longer than expected STR length. Precise and accurate detections of STR length are central for both basic and applied researches such as clinical diagnosis, genetic counseling, and forensic science. Various methods have been developed over the last 30 years to assay repeat length variations. The straightest methods are PCR-based techniques, such as fluorescence PCR, triplet-primed PCR, and small-pool PCR (Gymrek, 2017; Massey et al., 2018). However, stutter artifacts mediated by DNA polymerase slippage during PCR amplification are often introduced in a repetitive region (Kunkel, 1986; Walsh et al., 1996; Gymrek, 2017). Thus, it is difficult to distinguish between stutter artifacts and genuine STR polymorphisms.

In this study, we systematically investigated STR instability in human cells using GC-rich CAG and AT-rich ATTCT tandem repeats as examples. We found that triplicate bidirectional Sanger sequencing, PCR amplification under proper conditions, and next-generation sequencing (NGS) can accurately and reliably detect STR length. Finally, we found that STRs are intrinsically unstable in the human genome during cell culturing and single-cell cloning.

## Results

### Accurate detection of STR length by Sanger sequencing

Stutter noise could be generated by DNA polymerase slippage during Sanger sequencing. We first tested whether Sanger sequencing can generate accurate STR sequence traces. To this end, we separated one DNA sample into three replicates and performed Sanger sequencing with both forward and reverse primers (Fig. S1B and S1C). The AT-rich STR region of the *ATXN10* gene contains 16 ATTCT copy numbers in tandem (Fig. S1B). The signals of the forward sequence traces rapidly decline in or near the end region of STRs. However, signals of all three reverse traces have a consistent good quality (Fig. S1B). In addition, the GC-rich STR region of the *PPP2R2B* gene was bidirectionally sequenced in three replicates (Fig. S1C). All signals of both forward and reverse Sanger sequencing reads exhibit consistent sequence traces and good qualities (Fig. S1C). Therefore, for both AT-rich and GC-rich STRs, Sanger sequencing can generate consistent STR sequence traces with no stutter noises or overlapping peaks. We suggest bidirectional triplicate Sanger sequencing for accurate and reliable assaying of STR lengths.

### PCR conditions for faithful amplification of STR regions

We next evaluated stutter noises during PCR amplification. We first optimized PCR conditions to amplify a STR fragment with GC content higher than 65%. We used a super-fidelity DNA polymerase for excellent amplification of a high-GC content region. In addition, we extended pre-denaturation time from the usual 3 minutes to 6 minutes to fully denature the double-stranded genomic DNA. Moreover, we added 5% (v/v) dimethyl sulfoxide (DMSO) and 5% (v/v) glycerol to the PCR reaction mix. The former can bind to the major and minor grooves of the DNA template and inhibit its secondary structure, and the latter can help stabilize DNA polymerase activity (Sambrook et al., 1989). We performed three replicates of PCR experiments using genomic DNA of HEK293T cells for the *ATXN10* ATTCT and *PPP2R2B* CTG STRs. All PCR products display specific bands of the expected size on agarose gels with no “shadow” bands (Figs. 1A and S2A).

**Fig. 1.**
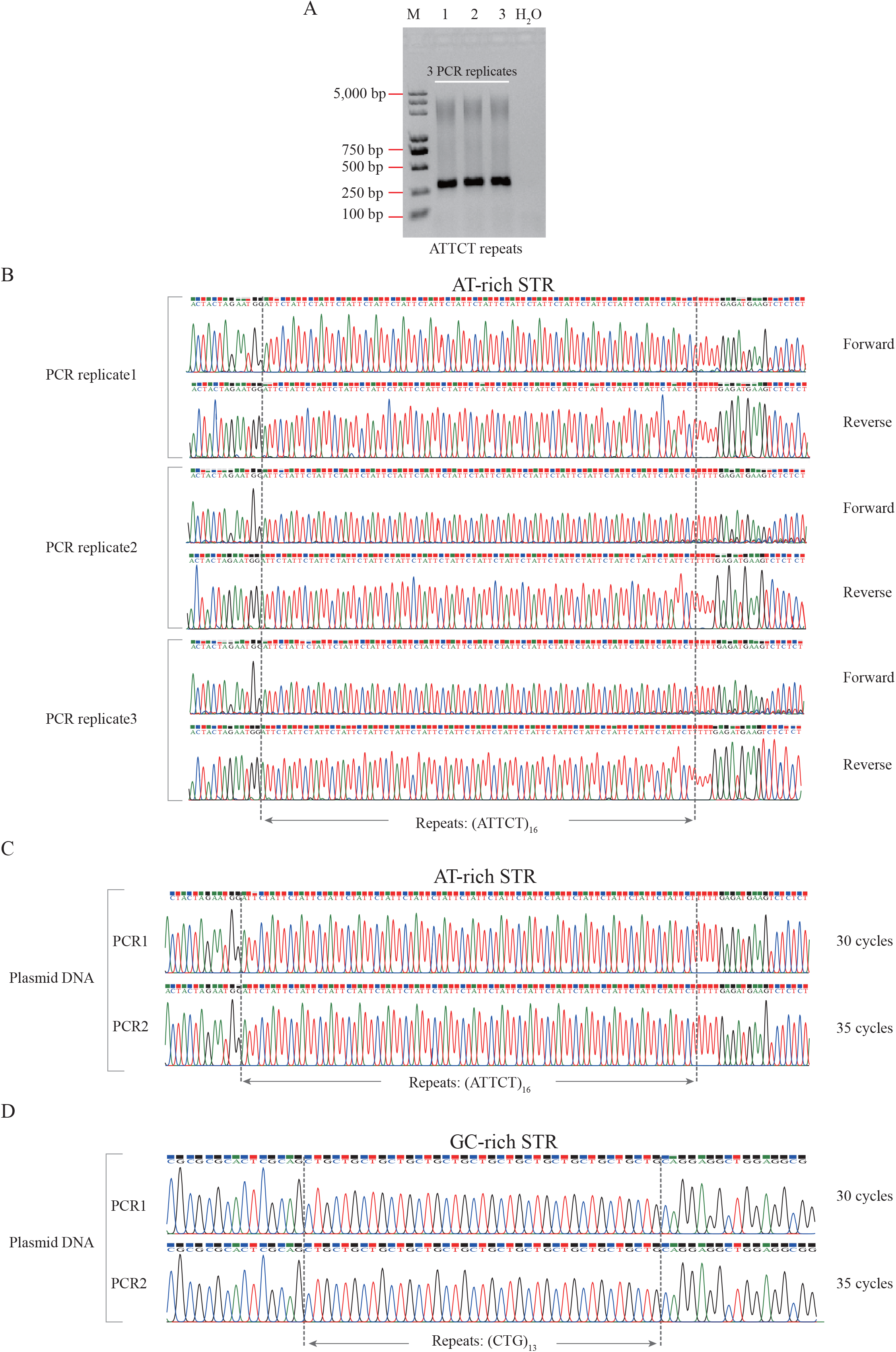
Evaluating stutter noises or artifacts of PCR amplification. **A:** An agarose gel electrophoretogram of PCR products containing ATTCT repeats. M: marker; Lanes 1, 2, and 3: three PCR replicates of the ATTCT STR in the *ATXN10* gene; H_2_O: negative control. **B:** Three PCR replicates of the ATTCT STR from a genomic template were sequenced with forward and reverse primers. **C** and **D:** Sanger sequencing of PCR amplifications from a plasmid template containing 16 ATTCT repeats (**C**) or 13 CAG repeats (**D**) with 30 or 35 cycles.

To further assess STR length accurately, all of the PCR amplicons were Sanger sequenced with both forward and reverse primers (Figs. 1B and S2B). Sixteen ATTCT repeats were observed with good trace signals in all of the genomic PCR replicates (Fig. 1B). However, one of three PCR replicates (replicate3) at 35 cycles for the GC-rich CTG STRs had overlapping Sanger traces near the 3’ terminal of tandem repeats (Fig. S2B). The overlapping signal traces indicated stutter errors of DNA polymerase slippage during PCR amplification of the genomic DNA template. We then tested different numbers of PCR cycles using the plasmid DNA as templates. We found that 30 or 35 cycles of PCR amplifications of plasmid templates resulted in good traces for the ATTCT (Fig. 1C) and CTG (Fig. 1D) STRs. Taken together, we concluded that triplicate PCR amplifications and Sanger sequencing can be used to precisely quantify STR length.

### STR instability in cultured human cell populations

To explore STR instability in cultured human cells, we freshly thawed a new vial of frozen HEK293T cells and cultured them for 19 days. After extracting the genomic DNA, ATTCT and CTG STR regions were amplified with 30 or 35 cycles and were sequenced. Surprisingly, sequencing data of both 30 and 35 cycles had overlapping trace signals at the 3’ terminal of STRs (Figs. 2A and S3A). To rule out potential stutter noises during PCR amplification, triplicate parallel Sanger sequencing was performed with both forward and reverse primers. Remarkably, we still observed overlapping peaks at the 3’ terminal of STR for all of the triplicate experiments (Figs. 2B and S3B). Compared with the initial genotyping of our HEK293T cells (Figs. 1, and S1-2), we concluded that 19-day human cell culturing can lead to STR length polymorphisms of the *ATXN10* and *PPP2R2B* genes. This suggests that the *ATXN10* and *PPP2R2B* STR regions were unstable in cell populations during human cell culturing.

**Fig. 2.**
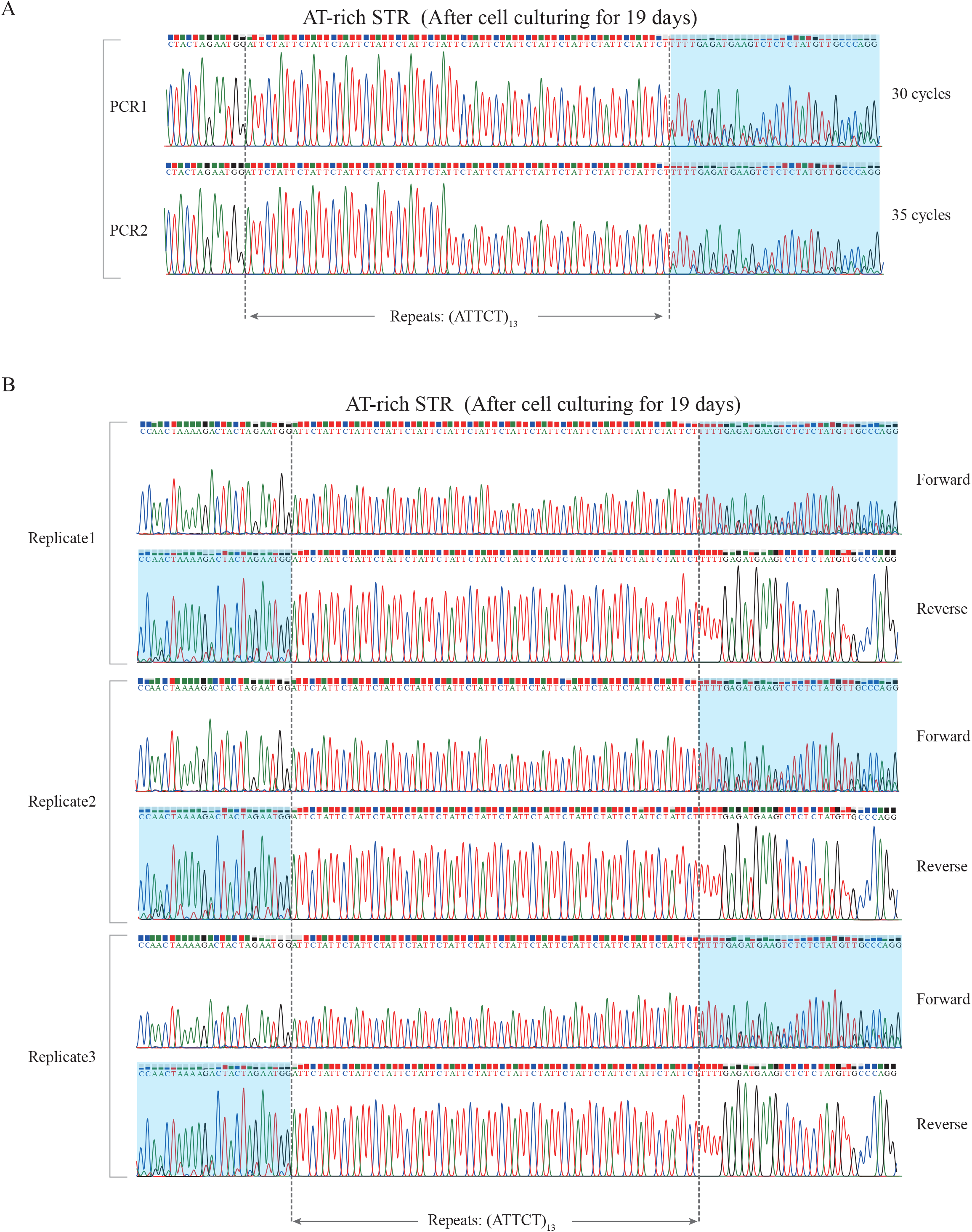
STR instability in cultured human cell populations. **A:** Overlapping trace signals were observed in human cell populations after 19-day culturing as detected by PCR with 30 or 35 cycles. **B:** Three replicates were sequenced with forward and reverse primers. The regions with overlapping trace signals were highlighted.

### Assessing STR length by next generation sequencing (NGS)

To determine the diversity of the mutated STR length in cultured human cell populations, NGS was applied to detect STR expansions and contractions. Sequencing libraries for the ATTCT and·CTG STR regions were prepared and sequenced on a NovaSeq platform. First, we evaluated the base quality of NGS reads and the Phred/Phrep scores of *PPP2R2B* and *ATXN10* samples were higher than 30 (Fig. 3A and 3B). By definition, this means that the probability of the incorrect base calling is less than 1 in 1000. Massive paired-end reads were merged (to ensure bidirectional sequencing of the STR regions), classified, sorted out, and quantified (Fig. 3C and 3D).

**Fig. 3.**
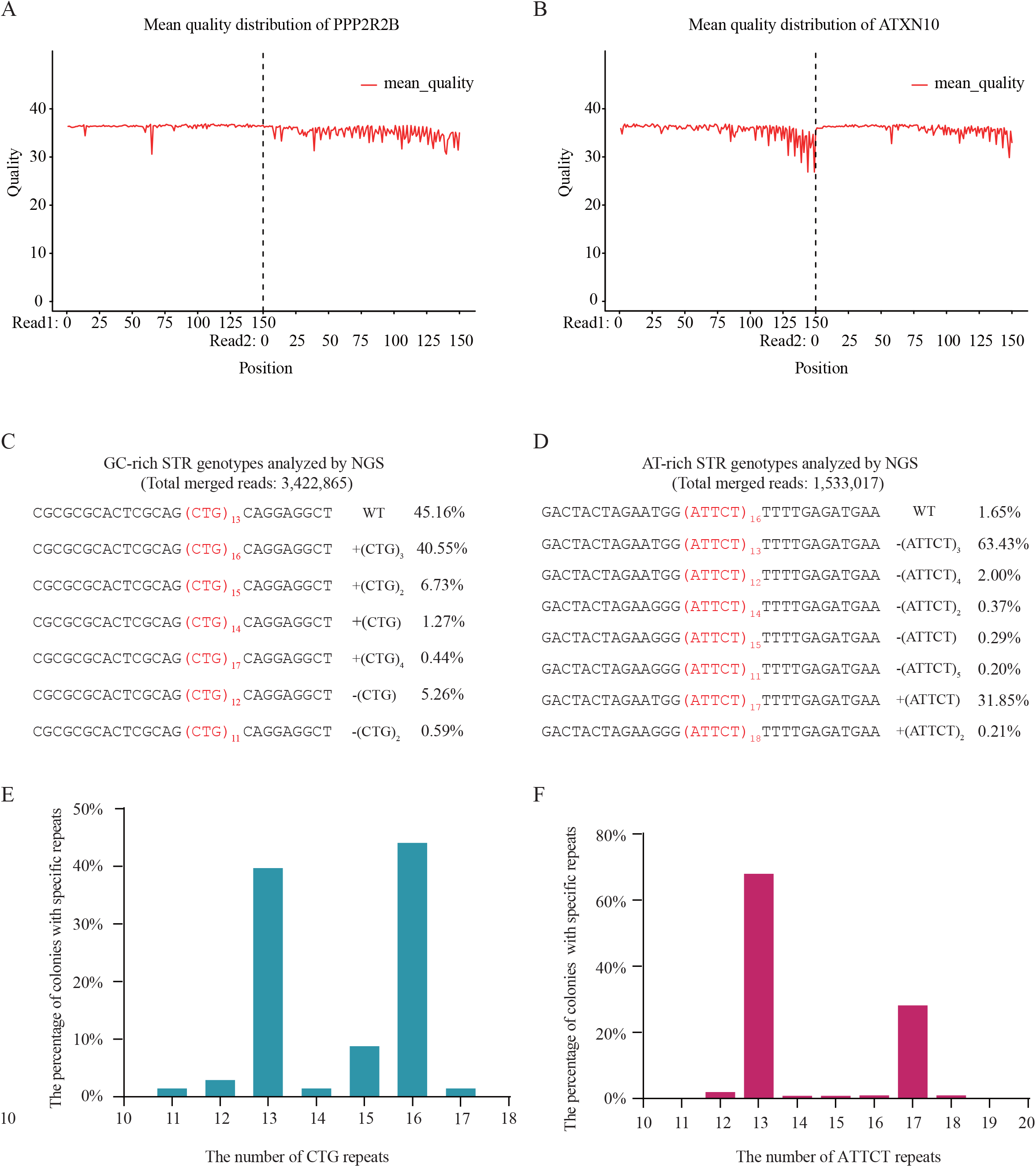
Genotyping STRs in 19-day cultured cell populations by NGS and Sanger sequencing. **A** and **B:** The Phred/Phrep scores of NGS reads for CTG repeats of the *PPP2R2B* gene (**A**) and ATTCT repeats of the *ATXN10* gene (**B**). **C** and **D:** NGS reveals the diversity of mutated STR length after 19-day culturing in cell populations from a STR containing 13 CTG repeats (**C**) or containing 16 ATTCT repeats (**D**). **E** and **F:** Confirmation of the NGS diversity of mutated CTG STR (**E**) and ATTCT STR (**F**) by bacterial cloning and Sanger sequencing.

For the GC-rich STRs, we observed that the majority (45.16%) remain to be the same as the wildtype with 13 CTG repeat units (Fig. 3C). In addition, we observed reads with the expansions of 1 to 4 repeats, which accounted for 48.99% in total. Among them, the expansion of 3 repeats occurred at a frequency of 40.55%. By contrast, contractions of 1 or 2 repeats occurred at a frequency of 5.85% in total (Fig. 3C). To confirm these NGS data for the CTG STRs, we cloned the Illumina library and randomly sequenced 68 clones by Sanger sequencing. The size and pattern of STRs from the Sanger sequencing data were largely consistent with the NGS data (Fig. 3E, comparing with Fig. 3C). Thus, the NGS approach could be used to detect the STR length polymorphism.

For the AT-rich STR of the ATTCT repeats, surprisingly we observed that the majority are not the wildtype STRs with 16 repeat units (Fig. 3D). In particular, we observed that the STRs with 13 repeat units are the most abundant merged reads which occupied 63.43%. In addition, the second abundant merged reads are STRs with 17 repeat units which occupied 31.85% (Fig. 3D). To confirm these NGS data of the ATTCT STRs, we randomly Sanger sequenced 103 clones from the Illumina library and observed consistent expansion or contraction ratios with the NGS data (Fig. 3F, comparing with Fig. 3D). Together, we conclude that the AT-rich STRs with ATTCT repeats are intrinsically more unstable than the GC-rich STRs with CTG repeats in cultured cell populations.

### STR instability in human single-cell clones

To further investigate the instability of STRs, we isolated single-cell clones by seeding an average of one cell into each well of the 96-well plates for continuing culture for 11 days (Fig. 4A). On the day of seeding cells, we harvested a sample of cell population and extracted their genomic DNA. STR genotyping was performed and the ATTCT or CTG STRs each had only one genotype (Figs. 4B and S4A). On the day 11, a total of 10 single-cell clones were genotyped. Remarkably, we observed that three of the ten clones (clones 3, 4, and 7) have overlapping trace signals for the ATTCT STRs (Fig. 4C-4E) and all of the other seven clones have single peaks, such as the clones 6 and 10 (Fig. S4B and S4C).

**Fig. 4.**
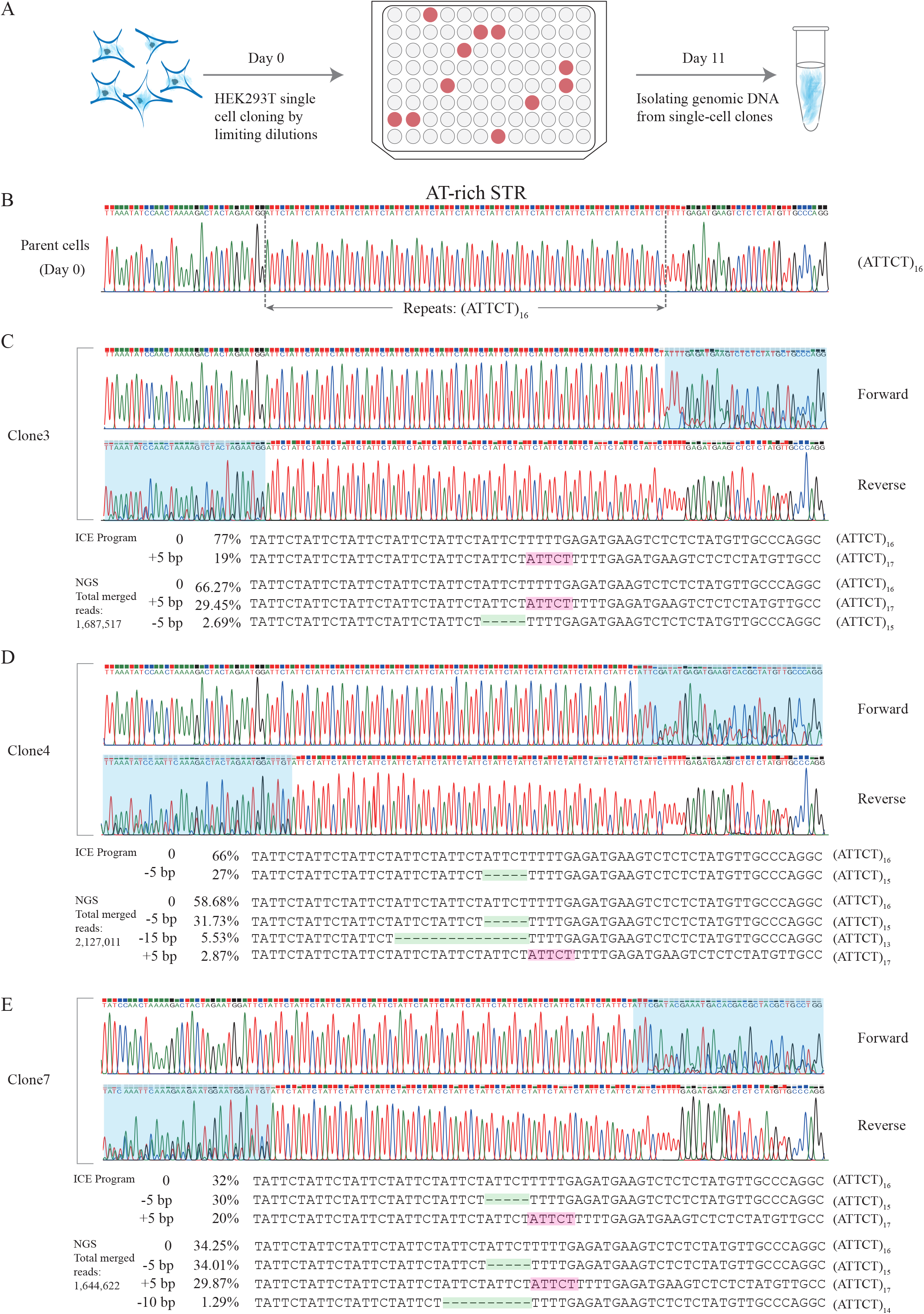
The instability of ATTCT repeats in human single-cell clones. **A:** Schematic of single-cell cloning from cultured human cells. **B:** Parent cells have only one repeat type. **C-E:** Diversity of mutated STR length in single-cell clones revealed by Sanger sequencing and NGS. All three clones have overlapping signals and their STRs have expansion or contraction.

We next analyzed the STR diversity in the three single-cell clones with overlapping traces by library construction and next generation sequencing (Fig. 4C-4E). Clone3 has an expansion of one repeat (29.45%) in addition to the repeat type of (ATTCT)_16_ (66.27%), while a contraction of one repeat only occupies 2.69% (Fig. 5C). Clone4 has the contractions of one repeat (31.73%) and three repeats (5.53%), and also has an expansion of one repeat (2.87%) (Fig. 4D). Moreover, Clone7 has both one-repeat contraction (34.01%) and expansion (29.87%), but a two-repeat contraction merely occupies 1.29% (Fig. 4E). We also used the ICE program (Inference of CRISPR Edits) (Conant et al., 2022) to analyze sequences for the three clones in the Sanger sequencing (Fig. 4C-4E). Comparing to the NGS data, the ICE data have the similar expansion or contraction patterns. Therefore, the ATTCT STRs have undergone copy number alterations during cell proliferation from single cells in the 11-day cell culturing. However, the CTG STRs of all of the ten single-cell clones display a single-repeat type of (CTG)_10_, as shown of Clone4 for an example (Fig. S4D). Their STR length is identical with the parent cell population (Fig. S4A), indicating that the CTG STRs are unaltered during the 11-day single-cell cloning. Therefore, the mutational rates of STRs are distinct for different STRs *in vivo*.

## Discussion

In this study, we systematically evaluated Sanger sequencing, PCR amplification, and NGS for assessing STR length. Although stutter noises occasionally exist, bidirectional triplicate Sanger sequencing with proper parallel PCR amplifications can precisely assay STR length. In addition, NGS with paired-end reads bidirectionally covering STR regions is a high-throughput and accurate sequencing method. Specifically, we systematically compared the NGS data with manual cloning followed by Sanger sequencing of the NGS libraries. We found that they are generally consistent. Finally, by assessing STR length via single-cell cloning of cultured human cells, we found that STRs are intrinsically unstable during human cell culturing. These data suggest that proper methods and keen awareness of stutter noises should be applied during assessing STR length in disease pathogenesis investigation, clinical diagnosis, and forensic science.

Although genomic repeat instability has been extensively studied in bacteria, yeast, mammalian cells, and patient tissues, the repeat expansion or contraction mechanisms still remain enigmatic (Kang et al., 1995; Maurer et al., 1996; Matsuura et al., 2004; Pelletier et al., 2005; Liu et al., 2007; Liu et al., 2010; Du et al., 2013). We carefully investigated STR instability during human cell culturing and single-cell cloning and found that STRs of the ATTCT repeats in the *ATXN10* gene or the CAG/CTG repeats in the *PPP2R2B* genes during the proliferation of HEK293T cells are intrinsically unstable. The CAG/CTG repeats are expanded during the 19 days of human cell culturing; whereas the ATTCT repeats are mainly contracted. For human single-cell clones, the instability is distinct for different tandem repeats. Specifically, three out of ten human single-cell clones have undergone expansion or contraction of the ATTCT repeat unit. By contrast, the CTG repeat traces of all of the ten human single-cell clones are unchanged, suggesting that the CTG repeats are more stable in human single-cell clones.

We observed an intriguing phenomenon that the forward sequencing signals for ATTCT repeats rapidly decline in the repeat regions; however, all of the reverse sequencing peaks still have a high quality (Fig. S1B). The decline of trace signals may be due to the formation of secondary structures in the forward sequencing template, which may partially block DNA polymerases. However, the reverse sequencing template does not form the polymerase-blocking secondary structures. We noted that this phenomenon occurred more frequently for the genomic DNA template than the plasmid DNA template. Interestingly, recent studies suggest that minidumbbell structures are formed by ATTCT pentanucleotide repeats (Guo and Lam, 2020). We suggest that the bidirectional triplicate Sanger sequencing should be performed in STR genotyping.

Once double strand breaks (DSBs) were generated in a tandem repeat region, it will increase the efficiency of tandem repeat instabilities (Wu and Shou, 2020). Specifically, DSBs induced by a programmed Zinc-finger or TALEN editors can lead to the contraction of the CAG/CTG STR (Mittelman et al., 2009; Richard et al., 2014). CRISPR technology developed in recent years has revolutionized genetic researches on genomic repetitive elements. In particular, recent studies revealed that the CRISPR/Cas9 system can generated staggered cleavages, resulting in 1 or 2 nucleotides insertions (Shou et al., 2018; Shi et al., 2019). It may be possible to engineer precise insertions utilizing Cas9 staggered cleavages to induce frameshift and to shorten STR length for future therapeutic development of repeat-expansion neurological diseases.

## Materials and methods

### Cell culture

The vials of HEK293T cells (human embryonic kidney cells, ATCC CRL-3216) were rapidly thawed by gentle agitation in a 37°C water bath (approximately 2 minutes). The freshly-thawed HEK293T cells were then cultured in Dulbecco’s modified Eagle’s medium (DMEM) (Gibco) containing 10% FBS (Gibco) and 1% penicillin-streptomycin (Invitrogen) in a cell culture incubator at 37°C with 5% (v/v) CO_2_. For cell preparations of single-cell cloning, the culture medium was firstly removed and discarded. The cells were rinsed by PBS to remove the FBS. Furthermore, the trypsin solution (Gibco) was added to the tissue-culture plate until cell layer was dispersed. The complete growth medium was then added into the plate. The cells were gently pipetted and centrifuged. Finally, the fresh medium was added to the pellet cells for cell suspension. The cells under good conditions were used to isolated single-cell clones.

### Genomic DNA extraction

Cells in a 6-well plate were harvested by centrifugation and washed with PBS once. The cells were then lysed by the Nuclei Lysis solution (Promega). Protein Precipitation Solution (Promega) was added to the lysed mixture, vortexed vigorously, and then centrifuged. The supernatant containing the DNA was carefully transferred to a clean 1.5 ml microcentrifuge tube. Cellular genomic DNA was then extracted by addition of room-temperature isopropanol. Finally the extracted genomic DNA was washed with 70% ethanol.

### PCR amplification

For GC-rich and AT-rich repeat fragments, super-fidelity DNA polymerase Phanta (Vazyme) for excellent amplification was used to amplify repetitive genomic regions. The PCR conditions were optimized by addition 5% (v/v) DMSO and 5% (v/v) glycerol. Furthermore, pre-denaturation time was extended from the usual 3 minutes to 6 minutes to fully separate the double-stranded genomic DNA. The PCR conditions are shown as below: pre-denaturing at 95°C for 6 min; followed by 35 or 30 cycles of 95°C denaturing for 15 sec, 60°C annealing for 15 sec, and 72°C extension for 50 sec; followed by a final extension at 72°C for 5 min.

### Plasmid construction

DNA fragments containing CAG or ATTCT STRs were amplified by PCR with specific primers (Table S1), and PCR products were then purified by PCR & DNA Cleanup Kit (NEB). STR PCR products were ligated with the pClone007 simple vector (Tsingke). Subsequently, the transformation was performed by using the *Stbl3* competent cells and the final positive colonies were inoculated in 2-ml LB medium and cultured at 37°C overnight for plasmid purification.

### Sanger sequencing and analyses

Purified PCR products or plasmids were sequenced by BigDye Terminator 3.1 (Applied Biosystems). Primers used for sequencing are listed in Table S1. For the overlapping Sanger traces, sequencing data were analyzed with the ICE Program (https://ice.synthego.com/#/).

### Single-cell cloning of human cell populations

Generation of human single-cell clones was performed as previously described (Li et al., 2015; Shou et al., 2018). Briefly, to isolate individual clones of HEK293T cells, cell populations under good conditions were seeded and cultured in 96-well plates at a density of one cell per well by limiting dilutions. Positive single-cell wells were identified on the day 6 and then cultured for additional 5 days. Cells in each well were collected and lysed by proteinase K (Sigma-Aldrich). Genomic DNA was then extracted and two microliters were used as PCR templates in a total volume of 25 µl.

### Next generation sequencing (NGS) and data analyses

To analyze multiple alleles of STRs, NGS libraries were prepared by a two-step PCR protocol. In the first step, STR DNA fragments were amplified using a pair of specific primers with 5’ terminal parts of P5 and P7 Illumina adapters (Table S1) and purified by 100% (v/v) SPRIselect beads (Beckman Coulter). In the second step, purified PCR products were used as PCR templates to generate the final NGS amplicon libraries with the barcoded P5 and P7 primers (Table S1). PCR libraries were then purified with 70% (v/v) SPRIselect beads and quantified by Qubit (Invitrogen). Finally, pooled barcoded libraries were sequenced on the NovaSeq platform (paired-end 150 bp). Paired-end reads of R1 and R2 were de-multiplexed by different index and assembled into one read using PANDAseq software (v2.8.1). For STR allele genotyping, each type of merged reads with different tandem repeat numbers was sorted out and calculated by a custom script. In total, we constructed 10 NGS libraries and obtained 48,509,956 raw reads covering each STR region bidirectionally.

## CRediT authorship contribution statement

**Yuzhe Liu** and **Jinhuan Li:** Investigation, Data curation, Validation, Writing - Original draft. **Qiang Wu:** Project administration, Resources, Supervision, Writing - Review & Editing.

## Conflict of interest

The authors declare no competing interests.

## Acknowledgments

This work was supported by National Key R&D Program of China (2022YFC3400200) and the Science and Technology Commission of Shanghai Municipality (19JC1412500 and 21DZ2210200).

## Figure legends

**Fig. S1.**
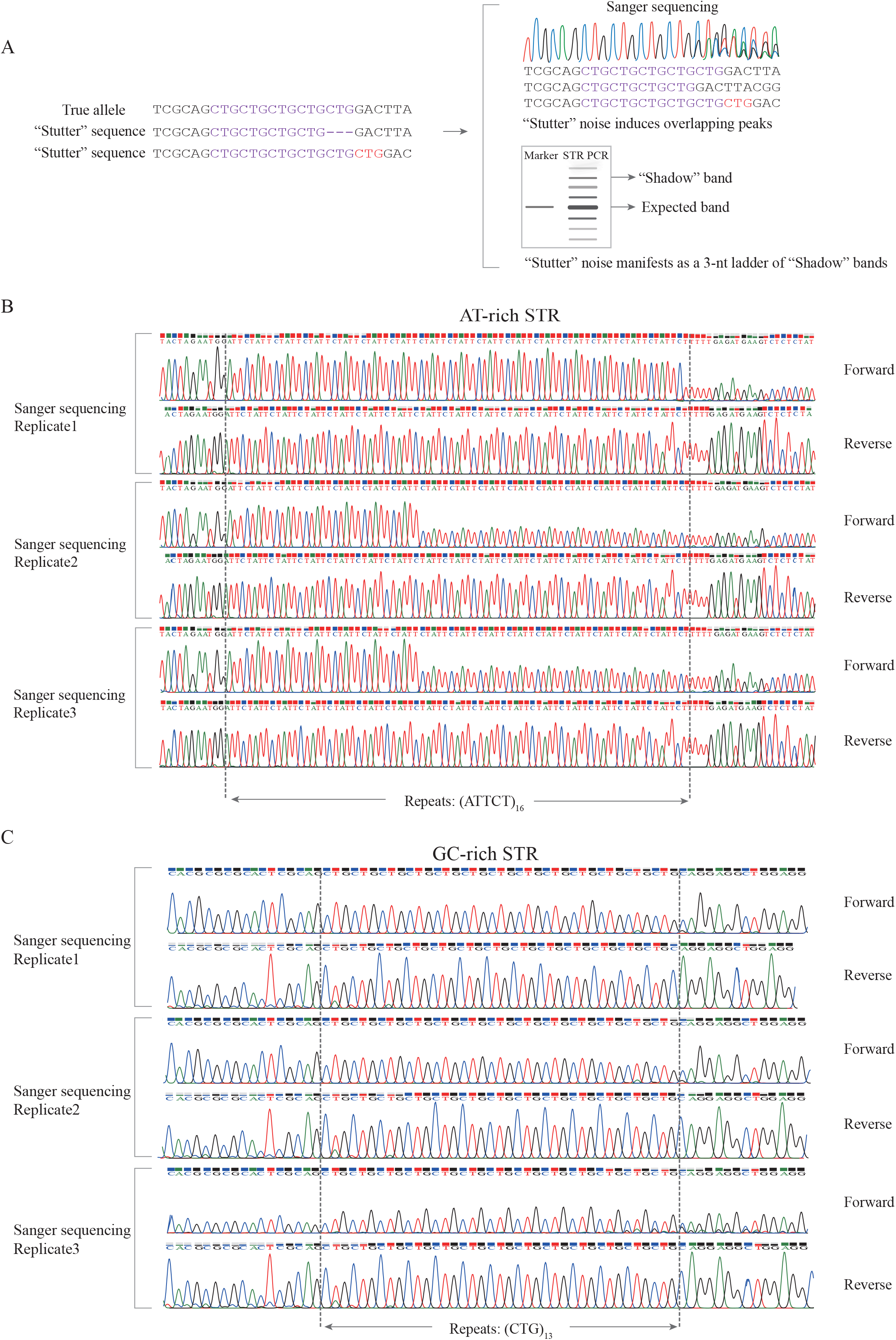
Accurate and reliable Sanger sequencing for STR detection. **A:** A schematic diagram introducing the stutter phenomenon. **B** and **C:** An AT-rich (**B**) or GC-rich (**C**) STR samples was divided into three replicates and Sanger sequenced with forward and reverse primers.

**Fig. S2.**
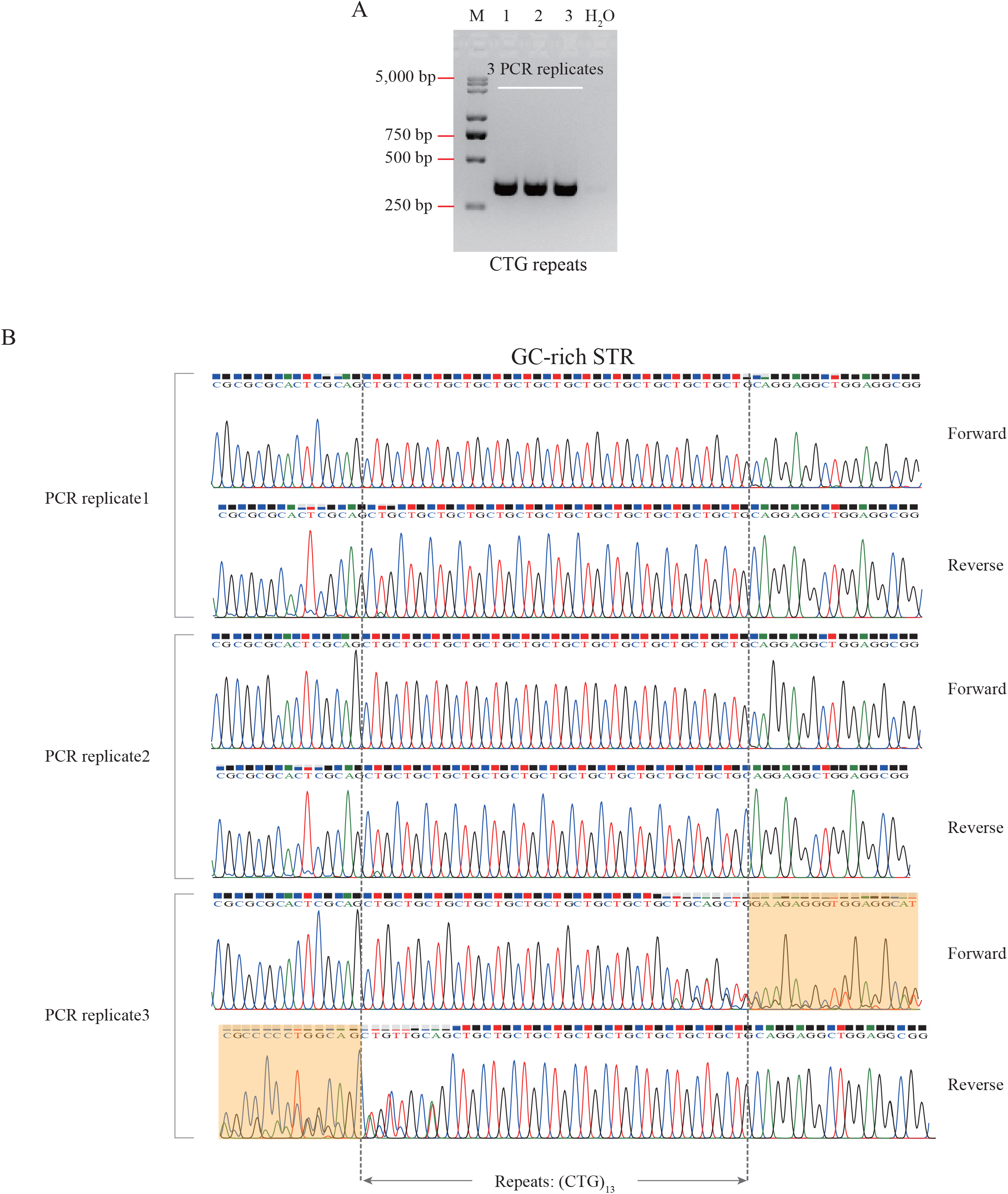
Evaluating stutter noises or artifacts of PCR amplification of CTG STR regions. **A:** An agarose gel electrophoretogram of PCR products with CTG repeats. M: marker; Lanes 1, 2, and 3: three PCR replicates of the CTG repeats in the *PPP2R2B* gene; H_2_O: negative control. **B:** Three PCR replicates of the CTG STR from a genomic template were sequenced with forward and reverse primers. Note that overlapping trace signals were observed in the third PCR replicate.

**Fig. S3.**
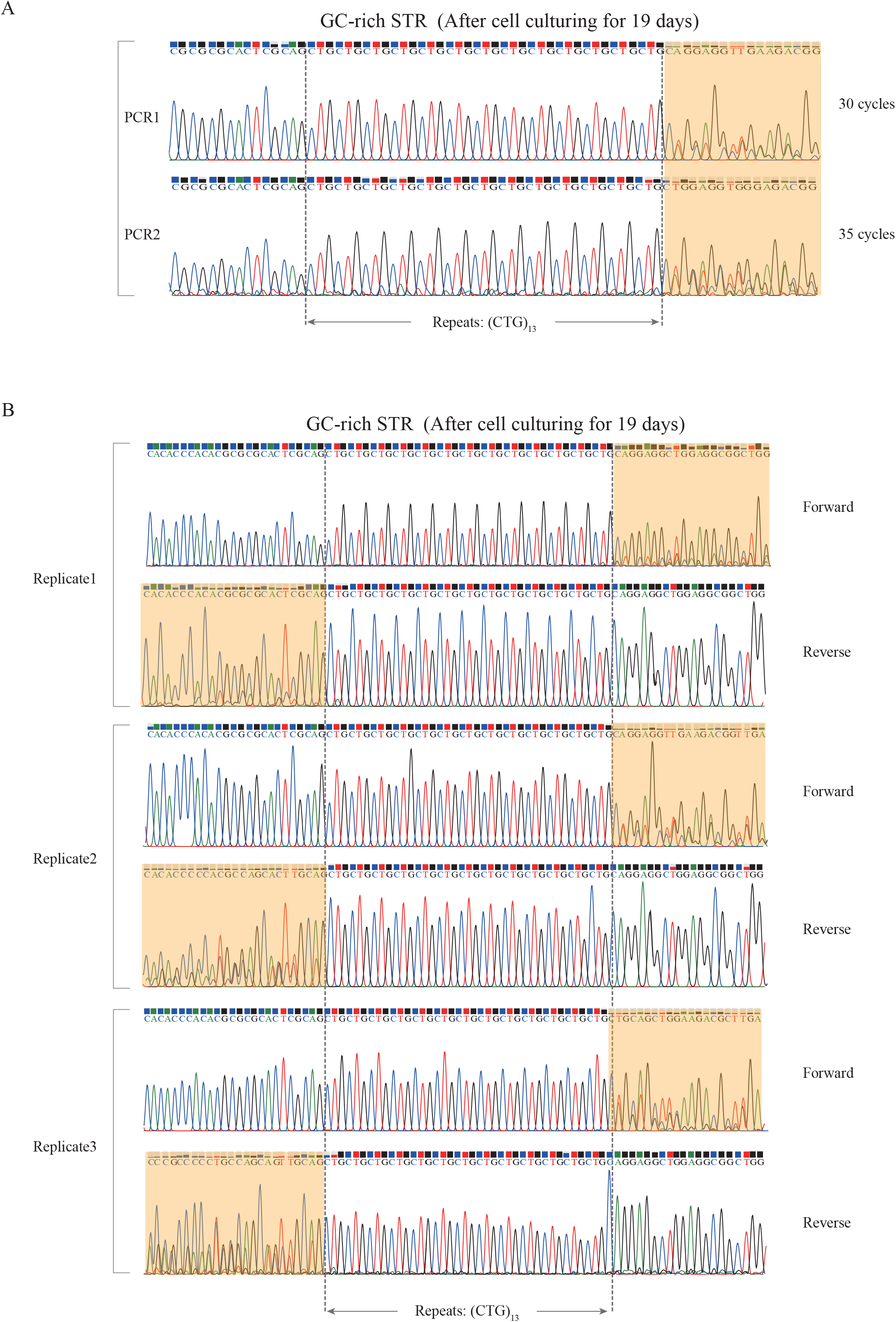
The instability of CTG repeats in cultured human cell populations. **A:** Overlapping trace signals were observed in human cell populations after 19-day culturing as detected by PCR with 30 or 35 cycles. **B:** Three replicates were sequenced with forward and reverse primers. The regions with overlapping trace signals were highlighted.

**Fig. S4.**
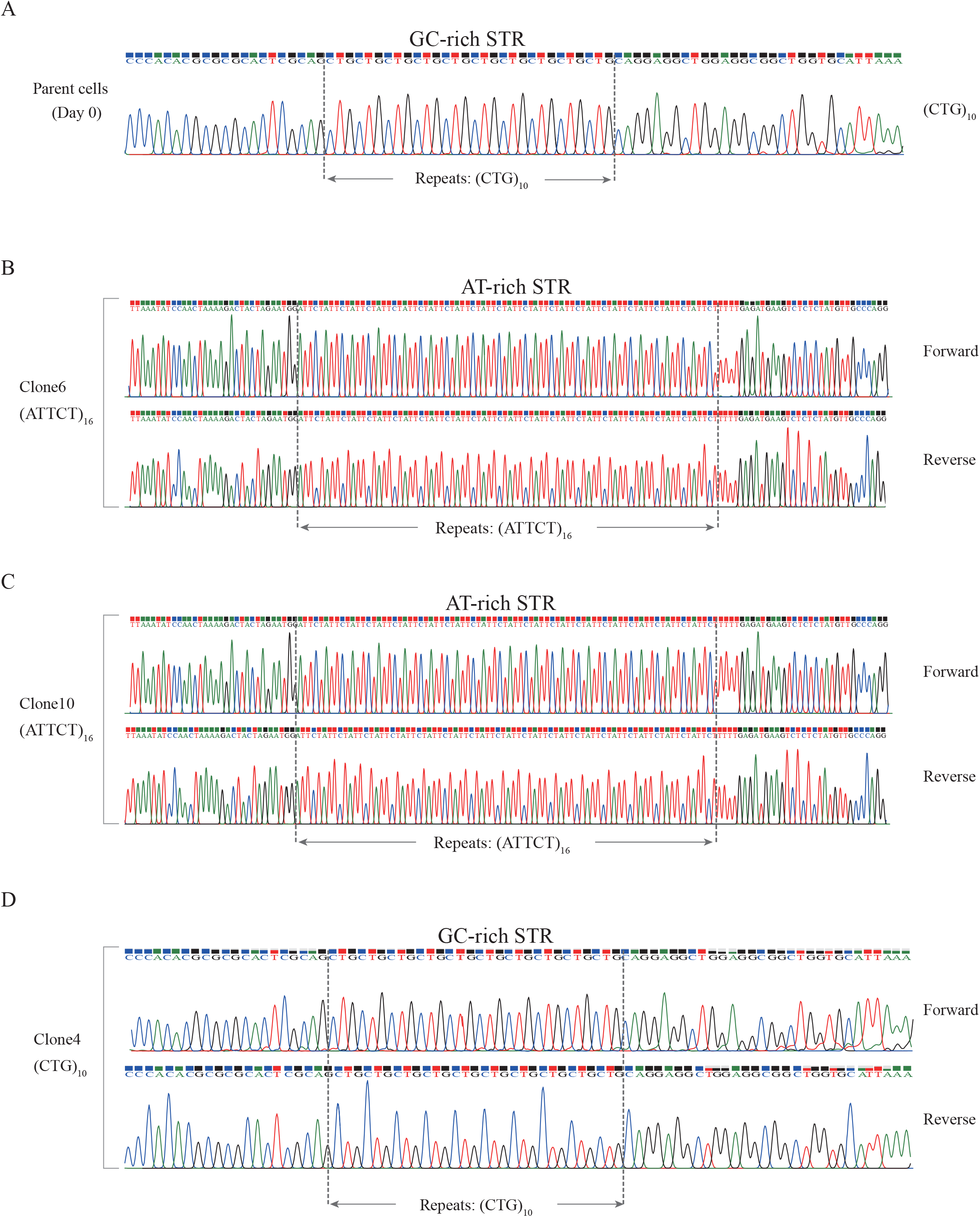
STR genotypes are identical with parent cells in some single-cell clones. **A:** Parent cells have only one repeat type in the CTG STR. **B-C:** The STR region contains 16 ATTCT repeats, which is identical with parent cells, in representative single-cell Clone6 and Clone10. **D:** The STR region contains 10 CTG repeats in representative single-cell Clone4, which is identical with parent cells.

**Table S1.**
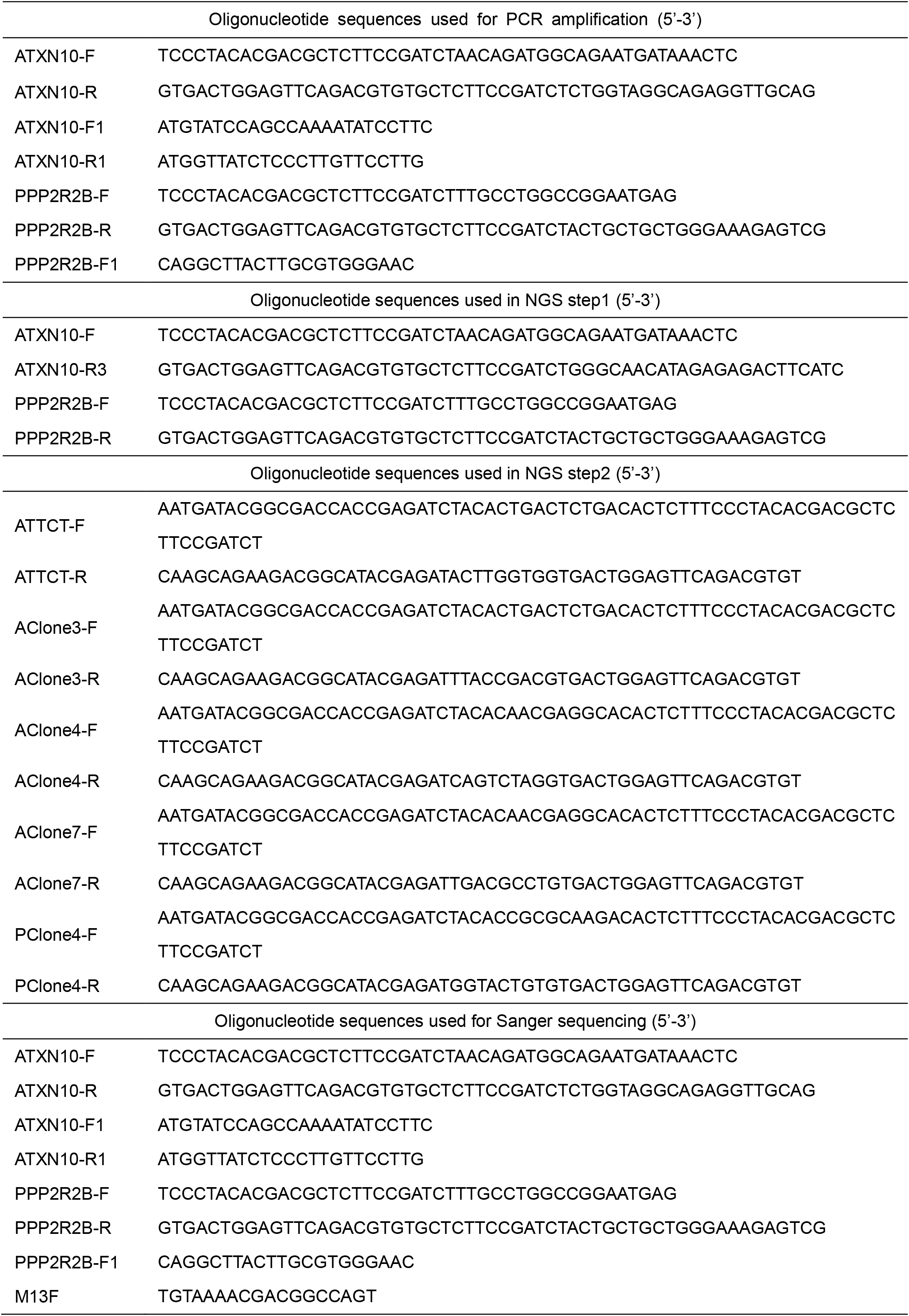
Oligonucleotide sequences used in this study.

## Notes

### Competing Interest Statement

The authors have declared no competing interest.

